# The Enterostat: a 3D-printed bioreactor for simulating gut microbiome dynamics

**DOI:** 10.1101/2025.08.21.671663

**Authors:** Emmi A. Mueller, Louis A. van der Elst, Alexander Gumennik, Jay T. Lennon

**Author notes:** Data accessibility: Code to reproduce all analyses is available on GitHub: http://github.com/LennonLab/Gut.

## Abstract

While the gut microbiome is central to host fitness, research in animal systems is limited by challenges in scalability and experimental control. Engineered bioreactors provide a valuable tool for studying microbiome dynamics, but many fail to capture the full anatomical and functional complexity of the gastrointestinal tract. Most existing models focus on either micro-scale features, such as villi and mucosal folds, or macro-scale parameters, such as fluid flow and volume, without integrating both. To address this gap, we introduce the Enterostat, a gut bioreactor platform designed to incorporate structural detail across scales while remaining adaptable to the dimensions and structural complexity of different hosts. We describe the modeling, fabrication, and operation of the Enterostat, and highlight its potential applications in microbiome research. The physical design of the Enterostat gives rise to oxygen gradients and particle retention, conditions that support the establishment and stability of microbial communities. The Enterostat also captures eco-evolutionary responses to antibiotic perturbations, demonstrating its utility for both applied and basic research. Overall, the Enterostat provides a versatile platform for pharmaceutical testing and for investigating how gastrointestinal architecture shapes microbiome composition and function.

## INTRODUCTION

The mammalian gastrointestinal (GI) tract harbors trillions of microorganisms, including bacteria, archaea, microeukaryotes, and viruses that are essential for host fitness (Lozupone et al. 2012; Costea et al. 2017; Gould et al. 2018). Collectively, the gut microbiome aids in digestion, supports immune system development, inhibits pathogen colonization, and contributes to the synthesis of essential vitamins (Sender et al. 2016; McKenney et al. 2018; Mueller et al. 2020). The composition and function of the gut microbiome is influenced by many factors, such as host diet, physiological stress, pharmaceutical treatments, and developmental stage (Turnbaugh et al. 2007; Costea et al. 2017; Rashidi et al. 2021; Dong et al. 2023). Disruptions to the gut microbiome have been linked to a range of health outcomes, including colorectal cancer, type 2 diabetes, obesity, inflammatory bowel disease, and anxiety (Cani 2018; Butler et al. 2023; Yadav et al. 2024; Fliegerová et al. 2025).

Understanding and managing the gut microbiome requires diverse experimental and analytical approaches that capture its complexity and functional dynamics. In clinical contexts, comparative studies and drug trials are considered the gold standard, but these methods are often constrained by ethical and logistical challenges (Vandeputte et al. 2021). Additionally, the limited use of invasive sampling make it difficult to identify the ecological or evolutionary processes driving changes in microbial communities (Vandeputte et al. 2017). Animal models offer a more tractable means of testing novel probiotics or dietary interventions, making them useful surrogates for human studies (Nguyen et al. 2015; Kumar and Atul 2024). However, they are expensive to maintain, and variation among individual animals can obscure experimental outcomes (Kumar and Atul 2024). In contrast, *in vitro* models provide a cost-effective and scalable alternative, especially when the system is easy to fabricate and operate (Zengler et al. 2019). These models are also attractive because they minimize host-associated variation and allow for controlled, replicated experiments without confounding factors that are inherent to animal models (Costa and Ahluwalia 2019; Roupar et al. 2021).

Accurate modeling of gut morphology and physiology is critical to the success of an *in vitro* system designed to study the gastrointestinal microbiome (Costa and Ahluwalia 2019). Morphological features can be broadly categorized based on the scale of anatomical structures. Macroscale features include the long, tubular geometry of the gut and the unidirectional flow created by peristaltic motion. Microscale features, such as villi, folds, and crypts, provide spatial organization that influence microbial colonization and interactions (Donaldson et al. 2016; Tropini et al. 2017). For example, villi in the small intestine are dense, typically 10-40 villi per mm^2^ (Standring 2015), and range from 0.5 to 1 mm in height (Hasan and Ferguson 1981). These physical features can influence fluid flow and retention (Fogler 2006; Wong et al. 2023), meaning that even in the absence of host cells, gut structure alone can shape microbial dynamics (Cremer et al. 2016; Müller et al. 2020). Physiological factors such as low oxygen levels, pH gradients, and the presence of a mucosal layer are also important characteristics of the gut environment (Donaldson et al. 2016).

To date, *in vitro* gut systems are generally divided into two main categories. The first includes large continuous or batch reactors that emphasize macroscale properties such as fluid dynamics and volume. For example, multi-stage systems like SHIME simulate different regions using connected vessels that vary in pH, operating volumes (0.3 L – 1.6 L), and dilution rates (Gibson et al. 1988; Molly et al. 1993; El Hage et al. 2019; Liu et al. 2025). These systems can operate stably for multiple weeks, but do not simulate shear stress or microstructural aspects of the gut (Venema and van den Abbeele 2013). The second category includes microfluidic devices, such as gut-on-a-chip models, which operate at very small volumes (< 1 mL) and focus on microscale features like tissue interfaces and flow-induced shear (Kim et al. 2012; Lee et al. 2024; Shin et al. 2024). Although these models provide fine spatial control, they are limited by short operational time frames, low microbial diversity, and flow regimes that poorly mimic gastrointestinal conditions (Wong et al. 2023). While both approaches have deepened our understanding mammalian gut microbiomes, a gap remains between them. Bridging it is important because flow rate, microstructure, and particle retention strongly interact in gut-like environments (Al-Mashhadani 2023).

Here, we introduce the Enterostat, a gut bioreactor system designed to combine both macro- and micro-scale features of the gastrointestinal tract into a single, flexible model. Using computer-aided design (CAD) and stereolithography (SLA) with biocompatibile materials, we developed an affordable and customizable 3D-printed bioreactor system that replicates the anatomical structure of a wide range of host gut environments. We describe the design and fabrication of the Enterostat and highlight its key morphological and biological features. A critical requirement of any gut model is the ability to reproduce biological responses to perturbation. We demonstrate this with a case study of antibiotic treatment evaluating microbiome stability and system response. We also discuss potential extensions of the Enterostat design, including adaptations for different host species, and explore how this platform can aid in a variety of applications, from testing nutritional additives and pharmaceuticals to investigating how gut structure influences microbiome dynamics.

## METHODS

We designed the Enterostat as a versatile bioreactor platform that can be readily adapted to mimic the anatomical complexity of gastrointestinal (GI) tracts across different host systems. Here, we describe the general features of the Enterostat design and present a human-gut-inspired prototype to illustrate its operation and potential applications. The system is built from a computer-aided design (CAD) model that captures key aspects of gut morphology while enabling customization of structural parameters to reflect different GI tracts. The physical model is fabricated using SLA 3D printing with photocrosslinkable resins. The wide variety of biocompatible and autoclavable resins available for use with SLA make it the most commonly used 3D printing method for biologically integrated devices such as prosthetics, dental fillings, and sealants (Kessler et al. 2020; van der Elst et al. 2020; Bayarsaikhan et al. 2021; Della Bona et al. 2021) and also makes SLA printing an ideal method for Enterostat fabrication. When running the Enterostat, fluid flow is managed by peristaltic pumps placed at either end of the model.

### Enterostat Design

#### Modeling

We began with a base model resembling a continuous-flow reactor and introduced surface complexity by adding folds and villi along the length of a tubular segment (Fig. 1). We developed our CAD model using Autodesk Inventor. In CAD modeling, the initial 2D sketches form the basis of the final 3D design. By defining geometrically critical features in these 2D sketches, subsequent extrusion and 3D operations can accurately represent the intended geometry, maintain dimensional relationships, allowing for a more precise translation from conceptual design to a manufacturable 3D model. Therefore, villi density and diameter were first patterned onto a 2D sketch (Fig. 1A), which was then projected onto the surface of a slice of the Enterostat (Fig. 1B). Villi were then extruded to the chosen height (Fig. 1C) and the resulting slice was used to generate a 1 mm-thick cross-section of the tube. These cross-sections were then stacked to produce segments of the desired length (Fig. 1D). Custom endcaps matching the tube’s diameter were designed with inlet ports sized to accommodate tubing used in experimental setups (Fig. 1E).

**Fig. 1.**
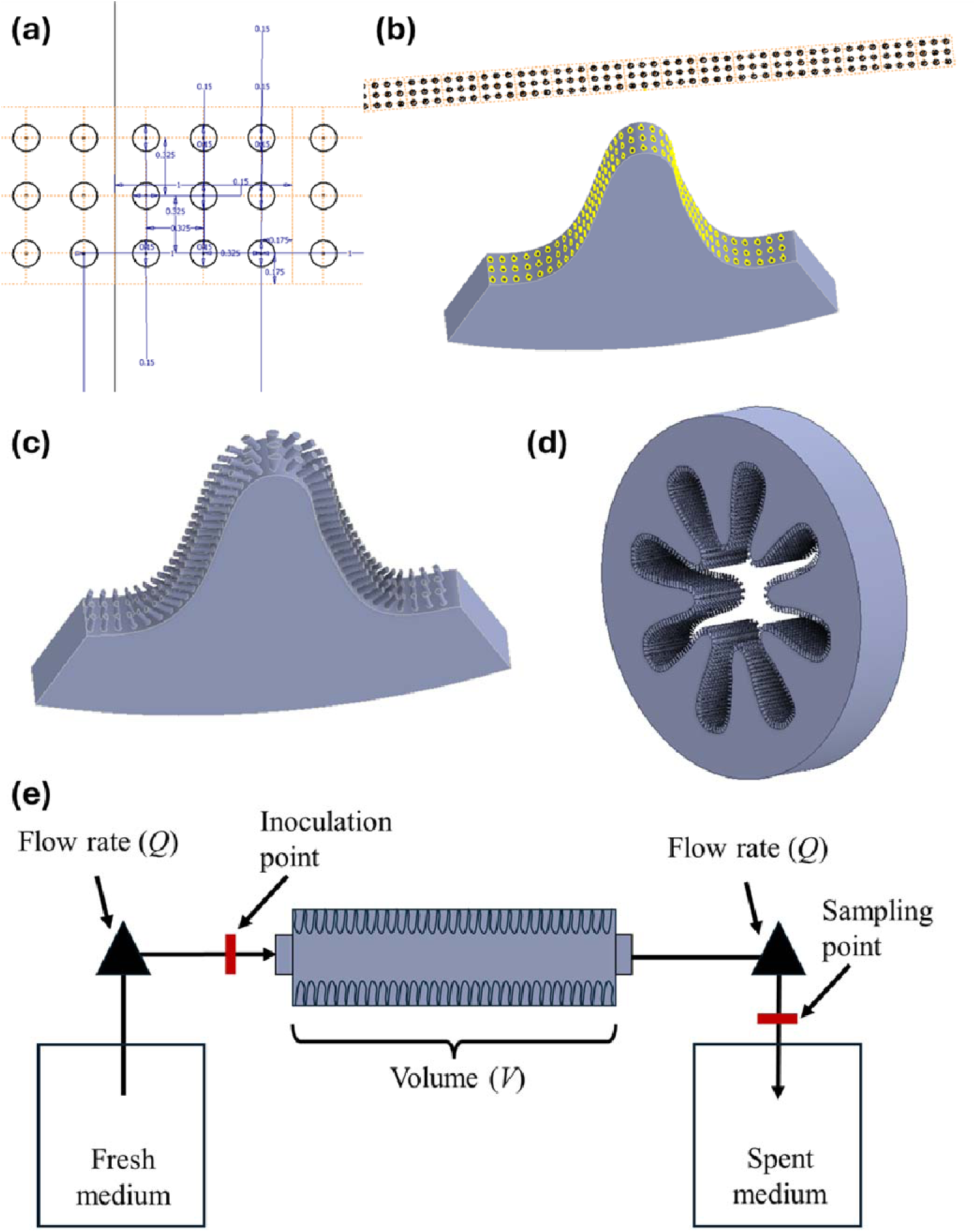
Enterostat design and operation. **(A)** Villi of a desired diameter and density are patterned onto a two-dimensional sketch. **(B)** This villi design is projected onto a three-dimensional section of the Enterostat cross-section. **(C)** Villi are extruded to the desired height and **(D)** the 1/8^th^ pie slice of the full cross section is patterned into a complete cross section and repeated to generate the desired total length of an Enterostat section. **(E)** After fabrication, Enterostats are connected to two peristaltic pumps using silicone tubing. The ends of the silicone tubing are attached to fresh medium and spent medium carboys. Inoculation point and sampling point are also indicated.

To demonstrate the flexibility and functionality of the Enterostat platform, we designed and fabricated a model based on the human ileum. The ileum is the distal section of the small intestine, typically measuring ∼3 m in length with an average internal diameter of ∼19 mm (Cronin et al. 2010; Standring 2015). Villi density in the small intestine ranges from 10 to 40 villi per mm^2^, with individual villus reaching heights of 0.5 to 1.0 mm (Standring 2015). In the ileum, villi are generally shorter and less densely packed compared to the more proximal regions of the small intestine (Standring 2015). Taking both anatomical features and the resolution limits of the Form2 3D printer (FormLabs, Sommerville, MA, USA) into account, we designed an ileum-inspired Enterostat version with a density of 9 villi per mm^2^. Each villus was modeled to have a diameter of 0.15 mm and height of 0.5 mm (Fig. 2A). The internal diameter of the gut segment varies from 8 mm to 22 mm along its length (Fig. 2C). The internal cavity is 95 mm long, accounting for approximately 3% of the total length of the human ileum (Fig. 2E). This version of the Enterostat has an internal working volume of 20 mL.

**Fig. 2.**
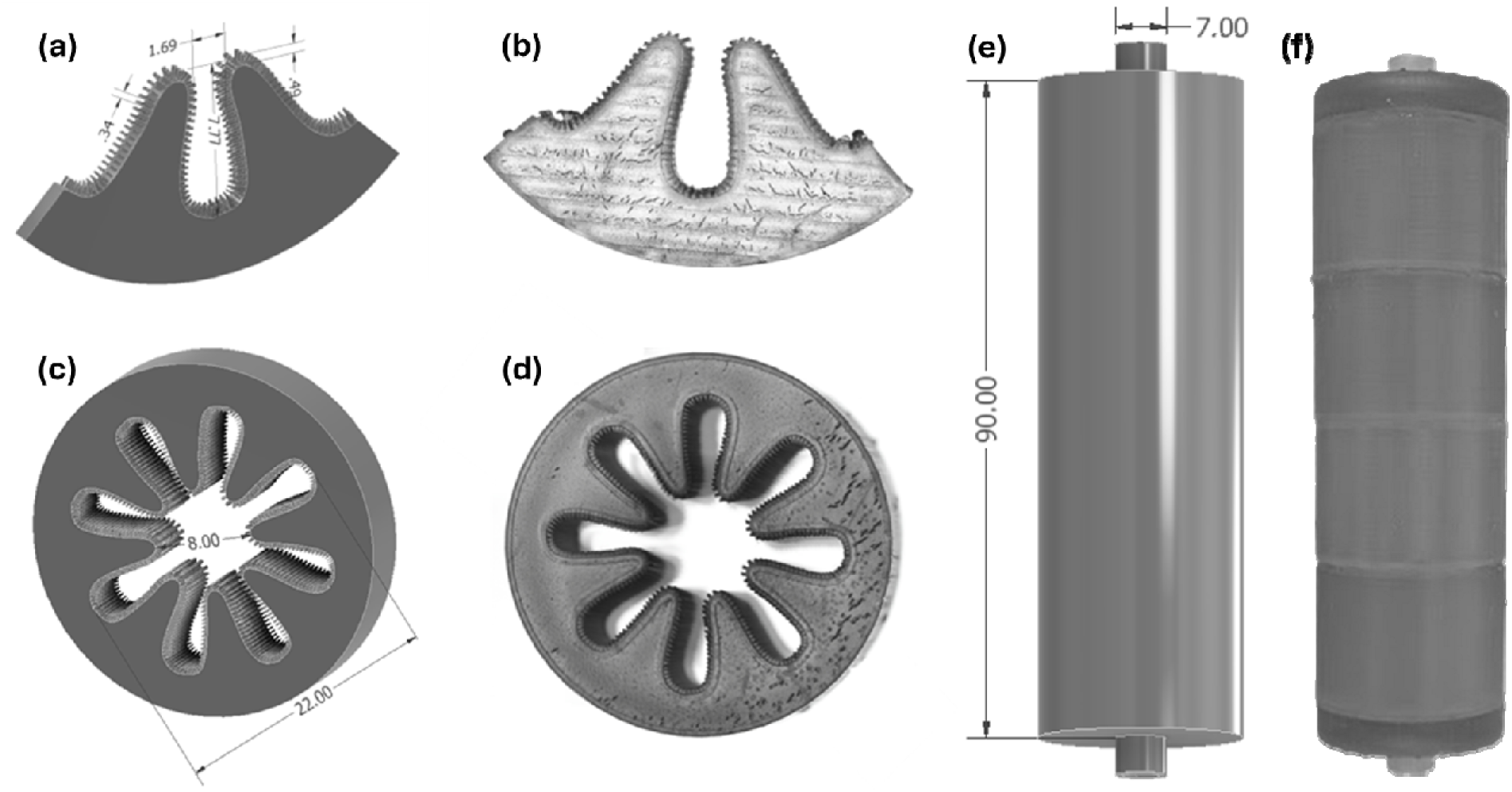
The Enterostat. (A) CAD model cutaway of the villi showing 1/8^th^ pie slice of the full cross section. Villi are spaced 0.34 mm apart and are 0.15 mm in diameter, generating a density of 9 villi per mm^2^. (B) The Enterostat was printed using FormLabs Surgical Guide resin and (C) the gut has an internal diameter of 22 mm at the widest point and 8 mm at the narrowest point in the folds. (D) The printed cross section shows villi separation along the folds of the Enterostat. The striation observed on flat surfaces in the printed Enterostats are due to the SLA print layering, but do not impact the bioreactor morphology. (E) The full length of the Enterostat is 90 mm, generating an internal volume (*V*) of 20 mL. It also has inlet and outlet connections that are 7 mm in diameter to allow tubing to be inserted and connected at each end. (F) Enterostat segments are attached and sealed with Loctite PL Marine Fast Cure Adhesive and allowed to cure for 24 h before being attached to silicone peristaltic pump tubing with heat shrink tubing.

#### Fabrication

We fabricated the Enterostat using SLA, a laser-based 3D-printing method in which a UV laser cures liquid resin layer by layer into a solid structure. SLA is well suited for applications requiring watertightness and fine surface detail owing to its high resolution, rapid print speed, and uniform bond formation across and between layers (Cosmi and Dal Maso 2020). Unlike fused deposition modeling (FDM), SLA avoids interlayer inconsistencies, making it ideal for constructing complex geometries such as villi. Additionally, SLA supports a wide range of resin materials, including biocompatible and autoclavable options developed for dental and medical uses (Turkyilmaz and Wilkins 2021).

To preserve fine structure, the Enterostat was printed in modular sections. The CAD model cavity segments were printed using Formlab’s Form2 SLA printer with Surgical Guide resin and an LT Resin Tank at a layer thickness of 0.05 mm (See Supplemental methods for Surgical Guide Resin specifications). After printing, parts were soaked in 99% isopropyl alcohol for 20 min with periodic agitation to remove uncured resin. Once dry, support structures were carefully removed, and the parts were post-cured using the Form Cure system (FormLabs, Sommerville, MA, USA) under the recommended conditions for Surgical Guide resin: 30 min of UV exposure at 60 °C. Gut segments were then assembled using Loctite PL Marine Fast Cure Adhesive and left to cure for a minimum of 24 h prior toautoclaving.

After autoclaving the assembled units at 121 °C and 15 psi for 30 min, we connected 1/8” diameter platinum-cured silicone tubing to the endcaps of the Enterostat using 1/4” heat-shrink tubing. To assess print fidelity, we compared the printed features of our human-ileum-inspired Enterostat to the original CAD specifications. Post-processed segments were stereoscopically imaged, and villus height and diameter were measured using ImageJ (*n* = 94). To quantify how closely the printed villi matched the modeled design, we calculated Cohen’s *d* between the specified villus dimensions and the distribution of measured dimensions from the printed Enterostat segments.

### Enterostat Operation

#### Set-up

After autoclaving, silicone tubing was attached to the Enterostat endcaps using heat shrink tubing to ensure a tight seal. Additional silicone tubing and polypropylene barbed fittings were used to connect the Enterostats to peristaltic pumps. Inflow and outflow reservoirs were connected to the silicone tubing at either end of the setup (Fig. 1E). To initiate operation, the first peristaltic pump was activated and set to the desired flow rate, allowing the Enterostat to fill completely. The second peristaltic pump was then started at the same flow rate, establishing a continuous, unidirectional flow through the system. This design helps prevent backflow and minimizes the risk of bacterial contamination in the inflow reservoir and tubing. During operation, the outflow reservoir was emptied and the inflow reservoir replaced with fresh medium as needed.

#### Particle retention

Given the unique physical structure of the Enterostat compared to commonly used chemostat-based gut models, we determined how the physical structure of the Enterostat alters flow through the system using particle tracing. We connected both styles of reactor to an inflow of water and peristaltic pumps which were operated over a flow rate gradient ranging from ∼1.5 to 17 mL/min. We used DayGlo Horizon Blue Paint as our tracer, injecting a 250 µL spike and then collecting one-minute fractions of outflow for 31 minutes. Each sample was homogenized and 200 µL was transferred to a 96-well plate for OD_600 nm_ measurement with an Epoch Microplate Spectrophotometer (Agilent, Santa Clara, CA, USA), along with a standard curve of DayGlo Horizon Blue Paint (OD_600 nm_). We calculated the percentage of particles remaining 30 min post-inoculation. Particle retention was analyzed using an indicator variable multiple regression (Lennon and Lehmkuhl 2016; Wisnoski et al. 2020), with flow rate as a continuous predictor variable and bioreactor type (i.e., chemostat vs. Enterostat) as a categorical variable. The regression includes an interaction term to assess whether the relationship between flow rate and particle retention differs between reactor types. Partial *R*^2^ values were generated using the ‘rsq.partial’ function from the ‘rsq’ package in R.

#### Operation

After assembly, we connected the gut models (*n* = 6) to multiflow Watson-Marlow 205U peristaltic pumps using 2-stop, peroxide-cured silicone tubing with a 2.79 mm internal diameter. For the inflow, we adapted a previously described gut medium (Molly et al. 1993), adding a potassium phosphate buffer to maintain a pH of 6 and omitting mucin (Table S1). Gut medium was supplied from a large carboy connected via 1/8” platinum-cured silicone tubing, and effluent from the Enterostats was collected in a separate carboy, which was emptied regularly throughout the experiment.

We inoculated the Enterostats with microbial communities derived from murine fecal pellets collected from male C57BL/6J mice. The pellets were stored at -80 °C in a 1:1 (vol:vol) mixture of Lysogeny Broth (LB) and 50% glycerol. Inoculation cultures from these fecal pellets were grown in LB overnight at 37 °C on a shaker table at 200 rotations per minute (rpm). At the start of the experiment, we added 1 mL of the overnight culture upstream of the Enterostat but downstream of the first peristaltic pump (Fig. 1E). We calibrated the flow rate of each peristaltic pump by converting rpm to mL/min, adjusting as needed to maintain matched inflow and outflow rates. For this operation test, pumps were set to a biologically relevant flow rate of 4 mL/min and run continuously for 7 d (Cremer et al. 2016).

#### Enterostat metabolism

To test whether the Enterostat’s structure could passively generate hypoxic conditions, we measured dissolved oxygen levels during benchtop operation and without external control. Using a PreSens SDR SensorDish Reader fit with PSt5 SensorVials (PreSens Precision Sensing GmbH, Regensburg, Germany), we filled the PreSens vials with 5 mL inflow gut medium and outflow from each gut in triplicate and sealed the vials. Ten minutes after lids were sealed, we measured dissolved oxygen (mg/L O_2_) with temperature set to 25 °C. PreSens vials were from batch #PSt5-0805-01. Differences in oxygen level between the gut inflow and outflow, were tested with a two-sample *t*-test (*n* = 152).

#### Reactor run-time and stability

To assess the stability of gut microbial communities in the Enterostats, we monitored bacterial abundance and community composition over a 7-d operational period. For bacterial abundance, we collected outflow from each gut and plated for total colony forming units (CFUs) on LB plates and antibiotic resistance abundances on LB plates supplemented with 20 µg/mL amoxicillin (stock solution of 15 mg/mL in DMSO, filter sterilized). We analyzed the temporal dynamics of total bacterial abundance with repeated measure analysis of variance (RM-ANOVA), using the ‘lme’ function from the ‘nmle’ package (version 3.1-164). Antibiotic treatment (control vs. antibiotic treated) was included as a between-subject fixed effect, and day as a within-subject fixed effect, with Enterostat specified as a random effect. Estimated marginal means were calculated using the ‘emmeans’ function from the ‘emmeans’ packages and letter ranks were generated using the ‘cld’ function from the ‘multcomp’ package (emmeans version 1.10.7; multcomp version 1.4-28). To quantify temporal community variability of the reactors, we calculated the coefficient of variance (CV) in total CFUs across time for each Enterostat (*n* = 12).

#### Microbiome diversity and compositional dynamics

To characterize microbial community dynamics in the Enterostats, we performed high-throughput 16S rRNA gene sequencing. Each day, 5 mL samples of effluent were centrifuged at 10,000 × *g* for 1 min. Supernatants were reduced to approximately 200 µL, and pellets were resuspended and stored at -80 °C for later processing. We extracted whole community DNA, prepared 16S rRNA gene libraries, and processed raw reads using the program mothur as described in greater detail (Locey et al. 2020; Wisnoski et al. 2020; Mueller and Lennon 2025, see Supplemental Methods). We used principal coordinates analysis (PCoA) to characterize changes in community composition both over time and with antibiotic treatment. For our PCoAs, we used the Jaccard dissimilarity metric from the ‘vegan’ package (version 2.6-4) in R (Oksanen et al. 2012).

### Enterostat Application

#### Antibiotic treatment

To demonstrate possible applications of the Enterostat and its capacity to capture gut-like biological dynamics, we tested the response of bioreactors (*n* = 6) to antibiotic exposure. Using the same experimental set-up as described above, after inoculation, we allowed the Enterostats to equilibrate for 4 d before treating these reactors with continuous antibiotics for the remaining 3 d of operation. We created the antibiotic treatment by adding amoxicillin to the medium at 2.5x the minimum inhibitory concentration (MIC) of our fecal pellet community (20 µg/mL; Fig. S1, See Supplemental Methods), from day 4 through the end of the experiment. To evaluate the effects of antibiotic exposure, we used a before-after-control-impact (BACI) design to capture both time-dependent changes in the Enterostat and treatment-specific responses (Underwood 1994).

#### Community response to antibiotics

To test the Enterostat response to antibiotics, we sampled gut outflow daily to observe total abundance, rates of antibiotic resistance, and changes in community composition both over time and with antibiotic treatment. Bacterial abundance and antibiotic resistance levels were determined as described above. Differences in resistant abundances with antibiotic treatment and time were analyzed using the same RM-ANOVA design as described above but day was replaced by timing (pre- and post-antibiotic exposure) as the within-subject fixed effect. To assess how community composition changes with the addition of antibiotics, we took daily samples and sequenced the 16S rRNA gene from the antibiotic-treated Enterostats and processed them as described above and in the Supplemental Methods. The effect of both time and antibiotic treatment on community composition was visualized using a PCoA as described above with the addition of ellipses representing 95% confidence intervals for the two treatments (‘geom_mark_ellipse’ function from the ‘ggforce’ package (version 0.4.2). Additionally, we conducted a PERMANOVA to test whether exposure to antibiotics altered community composition. We used the ‘how’ function from the ‘permute’ package (version 0.9-7) to block permutation by Enterostat ID and to account for the time series structure of the data.

To determine the identity of the resistant strains, we isolated antibiotic resistant colonies from amoxicillin-amended LB plates used for resistant abundance measurements. DNA extraction, PCR conditions, and sequencing preparation details can be found in the Supplemental Methods. Sanger sequencing was performed at Quintara Biosciences (Cambridge, MA, USA). We quality-trimmed the resulting sequences and generated BLAST results in Geneious Prime (Build 2025-05-19 14:11; Boston, MA, USA). All statistical analyses were performed in R (version 4.4.1; R Core Team 2024).

## RESULTS AND DISCUSSION

The Enterostat is a simple, flexible, reproducible *in vitro* gut model that captures both micro- and macro-scale features of the GI tract in a cost-effective and rapidly deployable format (Lennon et al. 2023). The unique 3D-printed design allows for easy modification to accommodate a range of host organisms and gut regions by adjusting features such as villus number and size, fold geometry, and segment length. Characterization of the Enterostat revealed physical structures that generated common abiotic conditions of the gut and supported stable microbial communities at biologically relevant flow rates for extended periods of time. Given these features, the Enterostat has potential applications in commercial testing of probiotics, nutritional additives, and drug delivery, and for fundamental research on how physical complexity shapes gut microbiome dynamics and host–microbe interactions.

### Enterostat Design

#### Modeling

We designed the Enterostat bioreactor to capture both micro- and macro-scale features of the GI tract while retaining flexibility to accommodate a wide range of experimental systems and questions. Leveraging Computer-Aided Design (CAD) software and Stereolithography (SLA) 3D printing technology, the system allows for unprecedented control over intricate anatomical and functional details. At the microscale, we successfully generated villi-like protrusions on folded surfaces (Fig. 2A,C), mimicking the complexity of the gut lumen. At the macroscale, these villi-bearing folds were assembled into a tubular structure that supported axial flow when connected to peristaltic pumps (Fig. 2E). Although the dimensions of the prototype were based on the human ileum, CAD modeling allows for straightforward modification of anatomical features for better biomimicry. For example, the length and diameter of the tube can be adjusted to match other regions of the GI tract, and villus density and height can be tuned to alter the degree of spatial complexity in the system (van der Elst 2024). This also allows the Enterostat to be used to model the GI tracts of a wide variety of hosts.

#### Fabrication and Operation

The fabricated human-ileum-inspired Enterostat successfully reproduced the key morphological features specified in the CAD design (Fig. 2). Fabrication of a single unit required 5 h of printing and 2 h for post-processing. The assembly using Loctite PL Marine Fast Cure Adhesive required an additional 24 h cure time. Thus, the total fabrication time was approximately two days, most of which involved passive steps such as printing and curing. Depending on the diameter and layout, multiple segments can be printed simultaneously, enabling parallel fabrication of several Enterostats.

Once assembled, the equipment required to operate the Enterostat is minimal, consisting of two peristaltic pumps, standard and 2-stop silicone tubing, and reservoirs for fresh and spent medium. While tubing and reservoirs can be cleaned and reused, the Enterostats and 2-stop tubing should be replaced between experiments. For our human-ileum-based Enterostat, the material cost per unit was approximately $30 USD, covering resin volume of assembled parts and connection tubing. The feasibility of running multiple Enterostats in parallel is primarily constrained by the medium consumption rate, with each Enterostat requiring roughly 6 L of medium per day, at a flow rate of 4 mL/min.

#### Printing precision

To evaluate how closely the printed structures matched the modeled dimensions, we compared the physical features of the fabricated Enterostat to the specifications of the CAD model, which was based on human ileum anatomy. In the design, villi were specified to be 0.5 mm in height and 0.15 mm in diameter. Using stereoscopic imaging, we measured 94 villi and found that the printed structures were, on average (± SEM), 0.430 ± 0.006 mm in height, slightly shorter than the modeled height (Cohen’s *d* = 1.11), and 0.251 ± 0.006 mm in diameter, slightly higher than the modeled diameter (Cohen’s *d* = 1.78; Fig. S2). Despite these modest deviations, the resulting dimensions remained within the range of villus sizes and densities typically observed in the human ileum (Standring 2015).

The current resolution of SLA printing, 25-50 µm, allows for accurate modeling of intricate anatomical details such as villi thickness, length, and density. However, as SLA printing technologies advance, the achievable resolution will continue to improve, enabling greater precision in replicating fine-scale gut anatomy (Yao et al. 2020). Additionally, the development of biocompatible resins for higher resolution methods such as digital light printing (Uchida and Bruschi 2024) will further enhance the fidelity of Enterostat structures. These improvements will be particularly important for modeling intestines of smaller hosts, such as those of mice, where villi average 0.22 mm in height and are below the current resolution of SLA (Gulbinowicz et al. 2004), as well as proximal regions of the human small intestine, which exhibit higher villus densities (Standring 2015).

### Enterostat Operation

#### Particle retention

The physical complexity built into the Enterostat, at both micro- and macro-scales, determines how organisms and resources move through the system and influences the abiotic conditions they experience (Fogler 2006; Nauman 2008; Cremer et al. 2016). For the Enterostat to function as an effective gut model, it must support the maintenance of microbial populations without allowing for complete washout. This can be achieved either through extended transit times that permit bacterial replication or through physical features that promote the physical retention of cells and particles even at higher flow rates (Mueller and Lennon 2025).

To evaluate how the Enterostat’s physical structure affects microbial retention, we compared its performance to that of a well-mixed, continuous-flow chemostat, across a range of biologically relevant flow rates (Cremer et al. 2016). Using indicator variable multiple regression, we found that particle retention in the Enterostat decreased linearly with increasing flow rate (slope = -0.027), dropping from ∼95% retention at the lowest flow rate (∼1.5 mL/min) to ∼50% at the highest flow rate (∼16.5 mL/min). Particle retention in the chemostat was unaffected by flow rate (slope = -0.002) because tracer particles were almost completely washed out within 30 min at al flow rates (Fig. 3, Table S2, *R*^2^ = 0.97, *F*_(3,12)_ = 139.9, *p* < 0.0001). While fast-growing bacterial species that can be found in GI tracts, such as *Escherichia coli*, *Klebsiella pneumoniae*, and *Salmonella enterica,* can replicate within this short window (Silva et al. 2009; Irwin et al. 2010; Liao et al. 2011), many others cannot (Weissman et al. 2021). Thus, our findings demonstrate that even under the highest gut-relevant flow rates, the Enterostat’s structural complexity promotes particle retention sufficient to support colonization, a key requirement for effective *in vitro* gut modeling.

**Fig. 3.**
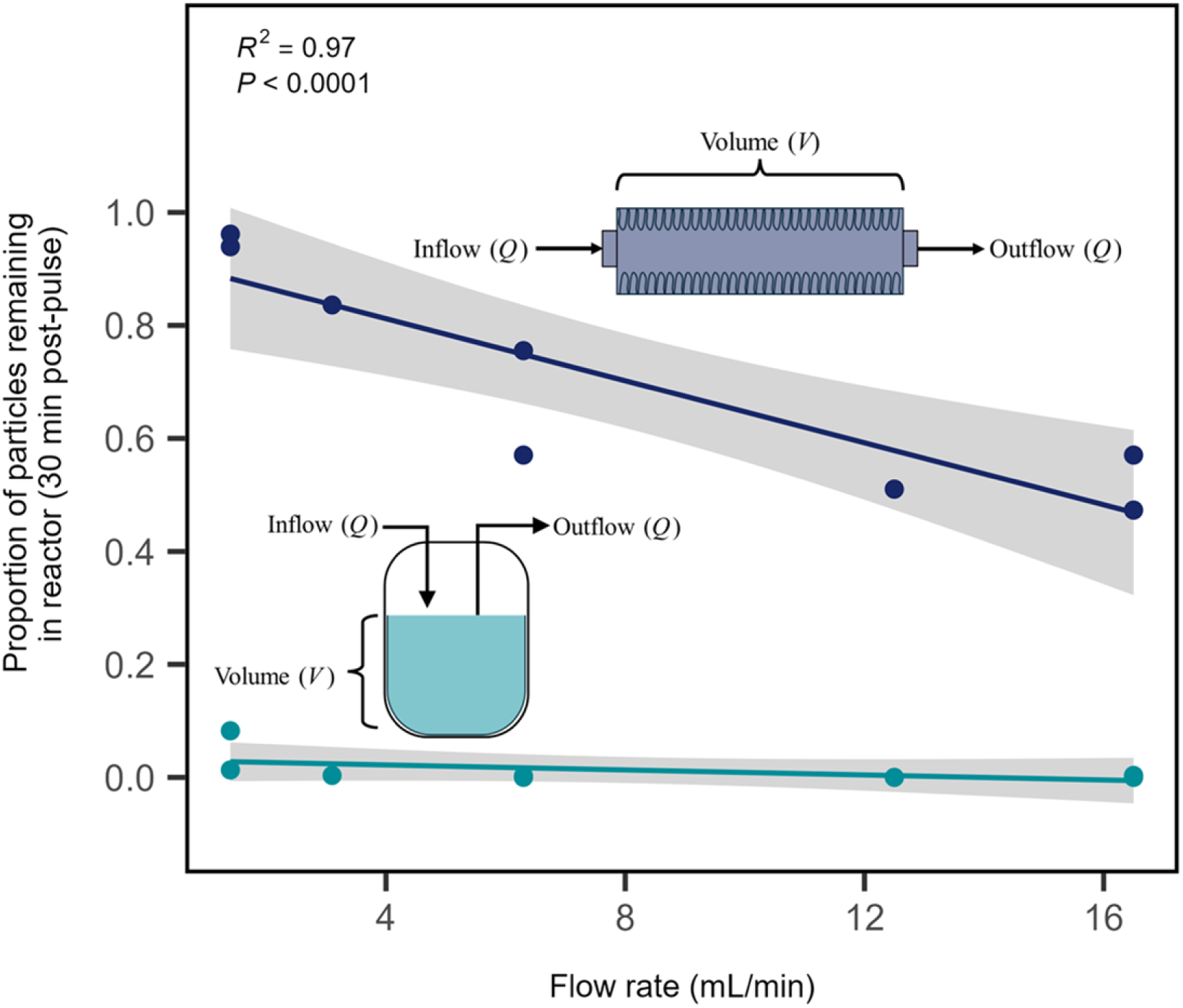
The Enterostat retains organisms and resources due to physical complexity. Particle tracing experiments show that at biologically relevant flow rates, a well-mixed chemostat experiences nearly complete washout after 30 min, regardless of flow rate. Comparatively, the rate of particle retention in an Enterostat of the same volume was flow rate dependent, with the highest flow rate, 16.5 mL min^-1^, retaining 47% of the pulsed particles, and the lowest flow rate, 1.5 mL/min, retaining 96% of the pulsed particles after 30 min.

#### Metabolism of the Enterostat

A key feature of the Enterostat is its ability to sustain hypoxic conditions without needing a glove box or other expensive gas-regulating apparatus. This property emerges naturally from its gut-inspired physical design, which mimics the geometry of the mammalian gastrointestinal tract. During operation, the Enterostat is filled with medium and perfused via peristaltic pumping, with no mixing. This configuration allows oxygen levels to decline over time, similar to conditions observed *in vivo* (Toerber et al. 1974). This enclosed structure, minimal headspace, and low gas diffusivity collectively generate steep oxygen gradients characteristic of the gut environment (Donaldson et al. 2016). Dissolved oxygen levels in the Enterostat outflow revealed hypoxic conditions (1.1 ± 0.1 mg/L O_2_) which were significantly lower than those in the fresh medium at the inflow (Fig. 4; *t*_141.18_ = 63, *p* < 0.0001). Notably, this reduction was achieved without any external control: the Enterostats were operated at room temperature on the benchtop under ambient atmospheric conditions. This passive reduction in oxygen is ideal for gut modeling, as modest increases in elevated oxygen availability (e.g.,1-5% atmospheric oxygen) are known to promote the proliferation of enteric pathogens (Wallace et al. 2016; Rivera-Chávez et al. 2017) and disrupt the stability of commensal microbial communities.

**Fig. 4.**
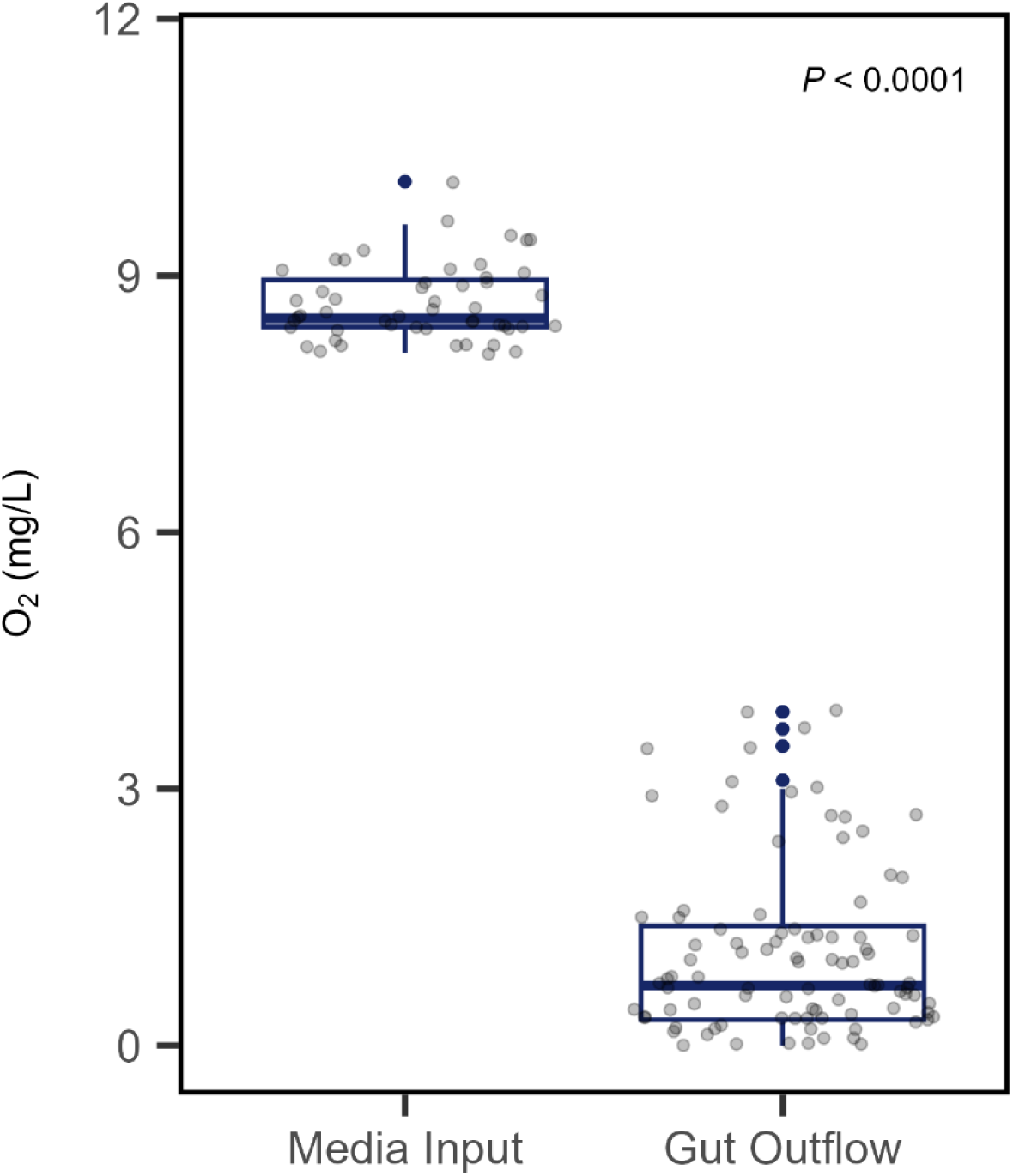
The Enterostat maintains hypoxic conditions reflective of the human ileum. The physical structure of the Enterostat generates hypoxic conditions without the need for an external low oxygen treatment. Medium entering the gut is continuously stirred, allowing for oxygenation while outflow from the gut has significantly lower oxygen levels.

### Enterostat Application

#### Biological stability

Stability of the gut microbiome is essential for host health and function (Fassarella et al. 2020). Therefore, the ability to maintain stable microbial communities, in particular, low temporal variation in abundances and composition, over extended periods is a requirement for any *in vitro* gut model. In our experiment, total bacterial abundance increased during the first 24 h following inoculation, likely reflecting an acclimation phase, and then remained relatively constant for the remainder of the one-week experiment (Fig. 5A, Table 1 & 2; Day, *F*_6_ = 9.37, *p* < 0.0001). After day 1, there were no significant differences in total abundance across time points. Moreover, the temporal variation in total abundance after the initial 24-h period was relatively low (CV = 85%), indicating that the Enterostat can operate in a consistent manner for extended periods of time, enabling investigation of microbiome dynamics and stability.

**Fig. 5.**
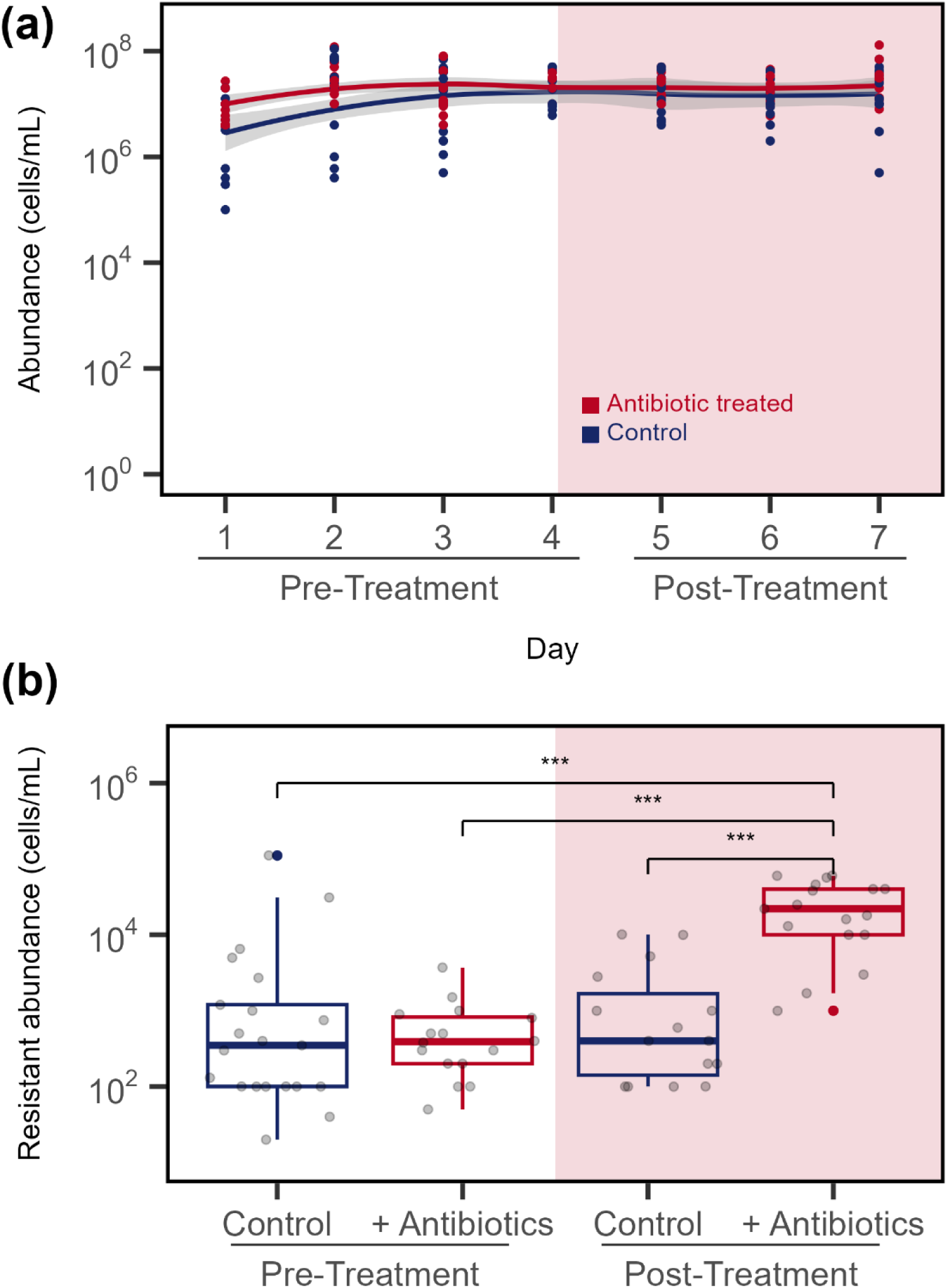
The Enterostat maintains steady state microbial communities, but also responds to perturbation. (A) After an initial increase in microbial abundance from day 1 to 2, microbial abundance remained stable at approximately 10^7^ cells/mL. A repeated measured analysis of variance (RM-ANOVA) revealed no significant changes either between days or between control and antibiotic treated abundances, meaning that the Enterostat is able to maintain a stable population size for at least one week after an initial acclimation period. While there was no change in total abundance with antibiotic treatment, (B) the rate of antibiotic resistance increased after addition of antibiotics compared to controls. The abundance of resistant bacteria was tested using a Before-After-Control-Impact (BACI) design where the interaction between treatment and timing indicates a significant increase in antibiotic resistance after antibiotic treatment began.

**Table 1.** Summary statistics for repeated-measures analysis of variance (RM-ANOVA) testing the effect of antibiotic exposure and timing on total abundance, resistant abundance, and observed taxonomic richness. Degrees of freedom (numerator DF & denominator DF) for each term (Source of variation) of the model are shown. Test statistics (*F*-value) are also shown. Significant relationships were determined from model *p*-values (α = 0.05).

| Model | Source of Variation | numerator DF | denominator DF | *F*-value | *p*-value |
| --- | --- | --- | --- | --- | --- |
| Total abundance | (Intercept) | 1 | 166 | 9641.03 | < 0.0001 |
|  | Day | 6 | 166 | 9.37 | < 0.0001 |
|  | ABX | 1 | 10 | 2.53 | 0.1429 |
|  | Day:ABX | 6 | 166 | 1.86 | 0.0909 |
| Resistant abundance | (Intercept) | 1 | 55 | 481.90 | < 0.0001 |
|  | Timing | 1 | 55 | 37.09 | < 0.0001 |
|  | ABX | 1 | 10 | 4.90 | 0.0513 |
|  | Timing:ABX | 1 | 55 | 26.14 | < 0.0001 |
| Observed richness | (Intercept) | 1 | 56 | 29.92 | < 0.0001 |
|  | Day | 6 | 56 | 1.65 | 0.1399 |
|  | ABX | 1 | 16 | 1.86 | 0.1918 |
|  | Day:ABX | 6 | 56 | 1.96 | 0.0769 |

**Table 2.** Summary statistics for total abundance and resistant abundance for the time periods and treatments. Estimated marginal means and standard error (EM means ± SE) are shown for each factor. Each Day or Timing:ABX interaction has a 95% confidence interval (95% CI) and a letter rank (Tukey rank) generated from a *post hoc* Tukey test.

|  | **Day** | **EM means ± SE** | **95% CI** | **Tukey rank** |
| --- | --- | --- | --- | --- |
| Total abundance<br>~ Day | 1 | 6.58 ± 0.112 | (6.33, 6.83) | a |
| | 2 | 7.30 ± 0.105 | (7.07, 7.53) | b |
| | 3 | 7.10 ± 0.101 | (6.88, 7.32) | b |
| | 4 | 7.39 ± 0.108 | (7.15, 7.63) | b |
| | 5 | 7.27 ± 0.102 | (7.04, 7.50) | b |
| | 6 | 7.19 ± 0.103 | (6.96, 7.42) | b |
| | 7 | 7.29 ± 0.100 | (7.07, 7.51) | b |
|  | **Timing:ABX** | **EM means ± SE** | **95% CI** | **Tukey rank** |
| Resistant<br>abundance ~<br>Timing * ABX | Pre- Control | 2.65 ± 0.214 | (2.18, 3.12) | a |
| | Pre- Antibiotics | 2.55 ± 0.226 | (2.05, 3.06) | a |
| | Post- Control | 2.81 ± 0.232 | (2.30, 3.32) | a |
| | Post- Antibiotics | 4.27 ± 0.226 | (3.76, 4.77) | b |

The ability to support a diverse microbial community is also an important feature of the Enterostat. Many *in vitro* models that focus on microscale gut features are only able to support a few microbial taxa (Marzorati et al. 2014; Shah et al. 2016; Lee et al. 2023). By comparison, over the 7 days, an Enterostat supported on average 32 (± 3.3) different bacterial taxa (97% 16S sequence similarity) and observed taxonomic richness was stable over time (Table 1; Day, *F*_(1,56)_ = 1.65, *p* = 0.1399). Across twelve Enterostats, global richness was 908 taxa. Compositionally, the Enterostats followed one of two trajectories, indicating some stochasticity during the acclimation period, However, community composition stabilized in all Enterostats by the third day (Fig. 6). Two days after initial establishment, a single *Bacillus* operational taxonomic unit (OTU) became numerically abundant across all Enterostats. This dominance was likely influenced in part by the overnight culture step, where rich medium favored certain fast-growing taxa (Fig. 6). The fact that this OTU was not initially dominant but rose to prominence over time suggests it was present in the inoculum but at low abundance. Other detected taxa included OTUs from the *Lactobacillales*, *Enterobacteriaceae*, *Staphylococcaceae*, and *Streptococcaceae* families, indicating that the Eneterostat provides a fabricated environment that is suitable for maintaining members typically found in a gut microbial community (The Human Microbiome Project Consortium 2012).

**Fig. 6.**
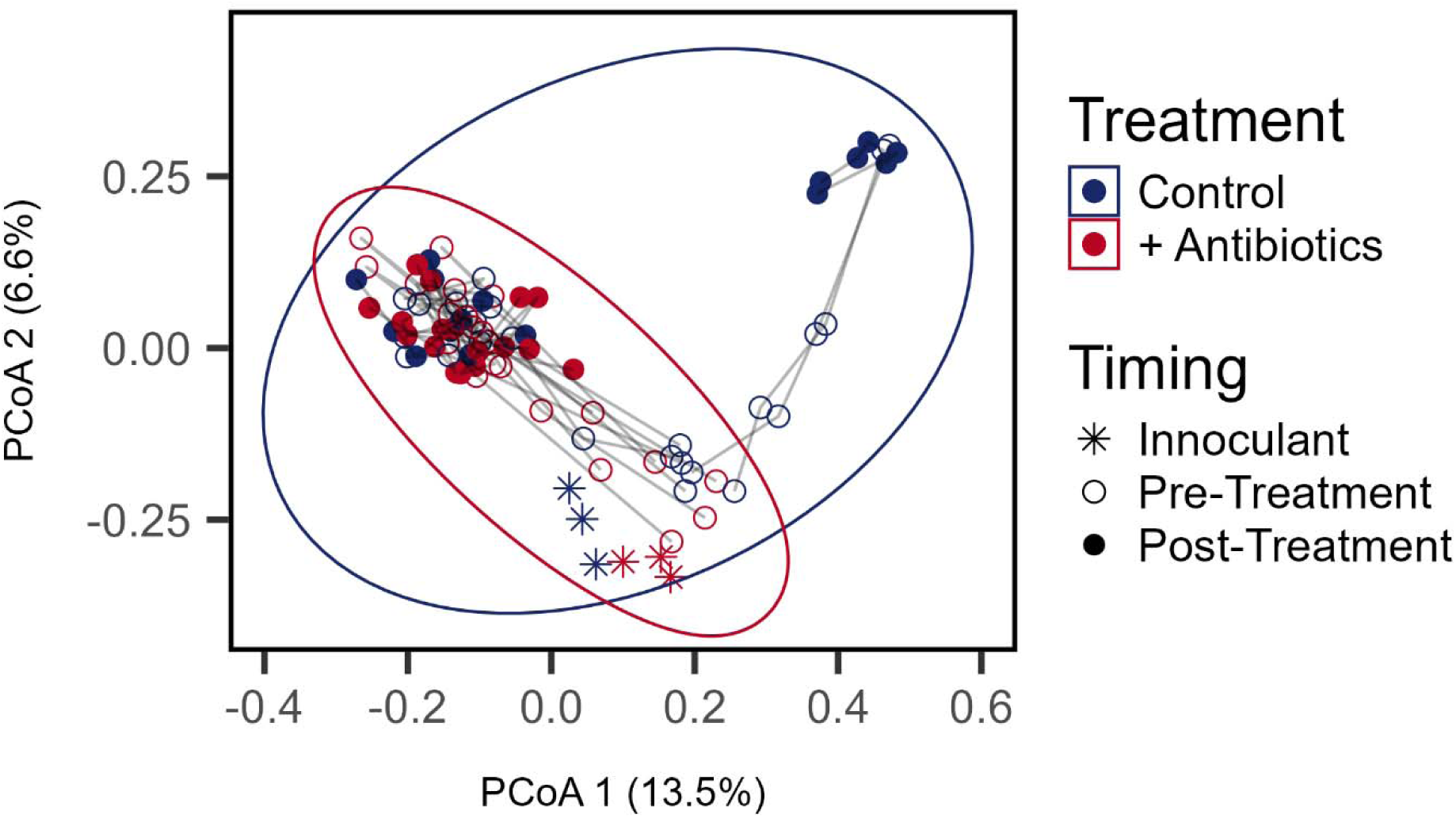
Microbial community composition reaches a steady state after two days. Principle coordinate analysis (PCoA) plot showing composition of the Enterostat microbial communities over the 7 days of operation. Distance between symbols represents dissimilarity between community compositions. Points are colored by treatment group with blue symbols representing Enterostats that never received antibiotics and red symbols representing Enterostats that received antibiotics on days 5 through 7. Stars represent the inoculating community. Open circles represent days 1 through 4 and closed circles represent days 5 through 7, when antibiotics were administered to the treated group. Ellipses represent 95% confidence intervals for the two treatment groups.

#### Evolution of antibiotic resistance

An effective *in vitro* gut model should be capable of capturing microbial responses to common perturbations, including antibiotic treatment. In our experiment, gut communities in the Enterostat remained stable following antibiotic exposure, with no changes in total bacterial abundance (Fig. 5A, Table 1; ABX, *F*_(1,10)_ =2.53, *p* = 0.1429), observed species richness (Table 1; ABX, *F*_(1,10)_ = 0.82, *p* = 0.3854), or community composition (Fig. 6; PERMANOVA, *F*_1_= 2.45, *p* = 1) compared to the no-drug control Enterostats. However, we observed a clear response of antibiotic-resistant microorganisms. Specifically, the absolute abundance of amoxicillin-resistant bacteria was significantly higher in drug-treated Enterostats compared to controls (Fig. 5B, Table 1; ABX:Timing, *F*_(1,55)_ = 26.14, *p* < 0.0001). Isolates taken from amoxicillin amended LB plates were overwhelmingly part of the order *Enterobacterales*, primarily strains of *Klebsiella oxytoca* and *Serratia liquefaciens* (Fig. S3). While we did not recover any *Klebsiella* among the OTUs generated with 16S rRNA sequencing, it is worth noting that on average, antibiotic resistant cells make up only 0.03% of the total abundance. If resistance is distributed across multiple rare OTUs, their low population sizes may place them below the detection threshold. We recommend that future studies increase sequencing depth or directly target the resistant community through selective enrichment or extraction of resistant organisms. Overall, the Enterostat’s ability to detect microbiome shifts in response to perturbation highlights its potential as a platform for studying clinically relevant processes, such as persister cell dynamics and the evolution of resistance under physiologically realistic gut conditions (Bakkeren et al. 2019; Baumgartner et al. 2020).

### Expanding Capabilities: Enterostat 2.0

Beyond the prototype presented here, the Enterostat platform is well positioned to address commercial needs and fundamental scientific questions, driving advancements in gut microbiome research. Its ability to reproduce intricate anatomical features and detect microbiome responses to perturbation makes it a promising tool for testing the effects of novel pharmaceuticals on gut microbial communities. Additionally, the Enterostat’s physical structure can be modified to model gastrointestinal disease states, including villus atrophy and surgical resection of gut segments, both of which are known to influence microbial composition (Fischer et al. 2017; Murray et al. 2017; Das et al. 2019). For example, decreased transit times due to ileostomy in patients with ulcerative colitis may lead to shifts in the microbial abundance or community structure (Tomita et al. 2004). Incorporating features such as a mucin layer or a colonic epithelial cell layer into the current model would enable investigation of host-microbiome interactions (Creff et al. 2019; Van Herreweghen et al. 2020). The design also supports real-time monitoring of biotic and abiotic conditions through the integration of embedded multi-material fiber devices (van der Elst et al. 2021). Fiber device integration can enhance Enterostat’s analytic functions by incorporating sensors, such as continuous pH monitoring or sonar-based biofilm monitoring using piezoelectric (PZT) elements (Faccini de Lima et al. 2019), as well enable fine-scale modulation of the cellular environment by precisely delivering biochemical agents directly to specific locations using porous fibers.

Looking forward, Enterostats may be constructed from living materials, transitioning beyond static representations to incorporate dynamic biological processes. The bioprinted hydrogel structures that are commonly used to model host tissues are often limited in size and mechanical stability due to their delicate nature. In future flexible versions of the Enterostat, fiber technology can also add tertiary structure and responsive actuation through embedded Shape Memory Alloy (SMA) fibers, which can be thermally induced to undergo controlled peristaltic-like contraction (Gokce et al. 2024; van der Elst 2024). As a more robust alternative, the internal cavity of the Enterostat could be coated with colonic epithelial cells to support studies of host-microbiome interactions (Murphy and Atala 2014). This comprehensive approach allows for the creation of sophisticated *ex vivo* organ models with microscale precision and real-time feedback, adaptable to host species of interest. Overall, the multi-scale engineering capability and inherent flexibility of the Enterostat, driven by digital manufacturing technologies, positions it as a valuable next step in the advancement of *in vitro* gut reactors microbiome research.

## ACKNOLWEDGEMENTS

We thank KJ Locey and BK Lehmkuhl for discussions about early design of the reactors. This research was supported by the National Science Foundation (DEB-1934554 and DBI-2022049 to JTL), US Army Research Office Grant (W911NF1410411, W911NF2210014 and W911NF2310054 to JTL) and the National Aeronautics and Space Administration (80NSSC20K0618 to JTL).

## Notes

http://github.com/LennonLab/Gut

### Competing Interest Statement

The authors have declared no competing interest.

