## supplement for "The Enterostat: a 3D-printed bioreactor for simulating gut microbiome dynamics"

### SUPPLEMENTAL METHODS

***Surgical Guide Resin specifications*** — Surgical Guide Resin contains 55-75% Urethane Dimethacrylate, 25-45% Methacrylate Monomer, and 1-2% Photoinitiator. Urethane Dimethacrylate is a common component in 3D printing resins, capable of favorable mechanical and biological properties when combined with other elements such as methacrylate monomers (Della Bona et al. 2021). Surgical Guide Resin is certified biocompatible per EN-ISO 10993-1:2009/AC:2010, confirming it is non-mutagenic, non-cytotoxic, non-sensitizing, and does not induce erythema, edema, or systemic toxicity.

***Community composition*** —We extracted genomic DNA using the DNeasy UltraClean Microbial Kit (Qiagen, Germantown, MA, USA). After suspending the cells in the kit provided PowerBead solution, but before the bead tube vortexing step, we incubated the samples with Proteinase K (10 µL 20 mg/mL) and Lysozyme (50 µL 50 mg/mL) for 1 h at 37 °C. After this initial step, we then proceeded according to the UltraClean kit instructions. DNA was eluted in 30 µL EB buffer and stored at -20 °C.

We performed PCR amplification of the V4 hypervariable region of the 16S rRNA gene using barcoded primers (515F and 806R) designed for the Illumina MiSeq platform (Caporaso et al. 2012). We used GoTaq DNA Polymerase (Promega, Madison, WI, USA) to run 50 µL reactions with the 5X Green GoTaq Reaction Buffer and 1 µL template DNA. The PCR reaction conditions were 98 °C for 30 s, 30 cycles of 98 °C for 10 s, 55 °C for 30 s, and 72 °C for 90 s, followed by a final extension of 72 °C for 5 min. We purified the sequence libraries using the AmPure Purification Kit (Beckman Coulter, Brea, CA, USA) and determined library concentration with the Quant-it PicoGreen dsDNA kit (Invitrogen, Waltham, MA, USA). We pooled the libraries to equal molar ratios (20 ng/library) and sequenced them using Illumina MiSeq 250 x 250 paired end reads (Reagents v2) at the Indiana University Center for Genomics and Bioinformatics.

***Diversity and assembly*** — We processed the raw 16S rRNA sequencing to determine microbial community composition stability in the Enterostat. We assembled the paired-end raw reads into contigs, quality-trimmed, and aligned them to the Silva database (Quast et al. 2013) before removing chimeric sequences using the VSEARCH algorithm (Rognes et al. 2016). We then split them based on RDP taxonomy (Cole et al. 2009) and binned sequences with 97% similarity using the OptiClust method (Westcott and Schloss 2017) to create operational taxonomic units (OTUs). We performed these initial sequencing processing steps using the software package mothur (Schloss et al. 2009; Schloss 2020). We removed samples with fewer than 10,000 reads and rarified the data to the sample with the fewest reads using the 'rrarefy' function ('vegan' version 2.6-10) in R (Oksanen et al. 2012).

***Minimum inhibitory concentration*** — To determine the MIC of the fecal pellet community we used to inoculate the Enterostats, we tested community growth over a gradient of antibiotic concentration. An overnight culture of fecal pellet community was added to a 96-well plate where each column contained a concentration of amoxicillin ranging from 0 μg/mL to 1,000 μg/mL. The plate was placed at 37 °C for 8 h, after which $OD_{600\ nm}$ was measured. We used a segmented regression to determine the breakpoint in the amoxicillin concentration ('segmented' function from the 'segmented' package; version 2.1-4). This resulted in an estimated MIC of 7.57 μg/mL (Fig. S1).

***Amoxicillin resistant isolates*** — Pure cultures of the isolates were extracted using the same method as whole community extractions described above. We performed PCR amplification of the full 16S rRNA gene (8F and 1492R) using Phusion High Fidelity DNA Polymerase (New England BioLabs, Ipswich, MA, USA) to run 50 µL reactions with the 5X Phusion HF buffer, 1.5 µL DMSO, and 1 µL template DNA. The conditions for the PCR reaction were 98 °C for 30 s, 30 cycles of 98 °C for 10 s, 58 °C for 30 s, 72 °C for 1:30 min, followed by a final extension at 72 °C for 10 min. We then purified the products using QIAquick PCR Purification Kit (Qiagen, Germantown, MA, USA), eluting in 30 µL of elution buffer. We then performed two BigDye Terminator (v. 3.1; Applied Biosystems, Waltham, MA, USA) reactions for each isolate, one for each of the two primers (8F and 1492R), using 1 µL BigDye MasterMix, 3 µL $MgCl_2$ , and 3 µL $H_2O$, with 1 µL template PCR product and 2 µL of primer.

### SUPPLEMENTAL TABLES

| Reagent | Amount per 1 L Medium | Unit |
|---|---|---|
| **Added before autoclaving** | | |
| Starch | 5 | g |
| Glucose | 0.4 | g |
| Yeast Extract | 3 | g |
| Proteose Peptone | 1 | g |
| Cysteine | 0.5 | g |
| NaHCO3 | 0.4 | g |
| NaCl | 0.08 | g |
| K2HPO4 | 2.09 | g |
| KH2PO4 | 11.975 | g |
| Tween 80 | 1 | mL |
| **Added after autoclaving** | | |
| CaCl2 | 0.008 | g |
| MgSO4 | 0.008 | g |
| Haemin | 0.005 | g |
| Trace Elements | 1 | mL |
| Vitamin solution | 1 | mL |
| **Vitamin solution (filter sterilized)** | | |
| Menadione | 1 | mg |
| Biotin | 2 | mg |
| Pantothenate | 10 | mg |
| Nicotinamide | 5 | mg |
| Vitamin B12 | 0.5 | mg |
| Thiamin | 4 | mg |
| Para-aminobenzoic Acid | 5 | mg |

**Table S1. Enterostat gut medium recipe**. This gut medium was adapted from a previously described gut medium (Molly et al. 1993), adding a potassium phosphate buffer to maintain a pH of 6 and omitting mucin.

| Response | Term | Estimate | SE | *t*-Statistic | *p*-value | Partial $R^2$ |
|---|---|---|---|---|---|---|
| Particle retention | Intercept | 0.922 | 0.042 | 21.89 | < 0.0001 | |
| | Flow rate | -0.027 | 0.004 | -6.49 | < 0.0001 | 0.00 |
| | Chemostat | -0.891 | 0.060 | -14.96 | < 0.0001 | 0.95 |
| | Flow rate x Chemostat | 0.025 | 0.006 | 4.221 | 0.0012 | 0.60 |

**Table S2. Summary statistics for relationship between flow rate and particle retention in a chemostat and the Enterostat.** Output from indicator variables multiple regression model testing how particle retention is altered by flow rate and bioreactor design. Model coefficients (Estimate) are shown for each term (Term) of the model. Significant linear relationships were determined from model *P*-values ($\alpha = 0.05$) and partial $R^2$ values were calculated. Test statistics (*t*-Statistic) and standard errors (SE) for each model term are also shown.

### SUPPLEMENTAL FIGURES

**Fig. S1. Minimum inhibitory concentration (MIC) of inoculating microbial community.** Amoxicillin was diluted to concentrations ranging from 1 mg/mL to 1 µg/mL and inoculated with the same microbial community used to inoculate the Enterostats. Based on a segmented regression, the minimum inhibitory concentration of the community in amoxicillin is ~8 ug/mL. As resistance tends to occur at 2 to 4 times the MIC, we chose a dose of 20 µg/mL as our treatment, approximately 2.5x the MIC, to ensure that the communities would react to the concentration without wiping out the entire community.

**Fig. S2. Enterostat villi printing resolution.** To model the human ileum, the Enterostat CAD model printed for experiments had a villus height of 0.5 mm and a villus diameter of 0.15 mm. The (A) villus height and (B) villus diameter of the printed Enterostat were measured and distributions were plotted with vertical dashed lines representing the CAD model values.

**Fig. S3. Heatmap of 16S rRNA gene sequence similarity between amoxicillin resistant isolates.** All isolates, aside from AMXI_30, were taxa from the order *Enterobacterales*, primarily *Klebsiella oxytoca,* which make up the larger high similarity cluster, and *Serratia* l*iquefaciens,* which make up the smaller high similarity cluster.

**Fig. S1.**

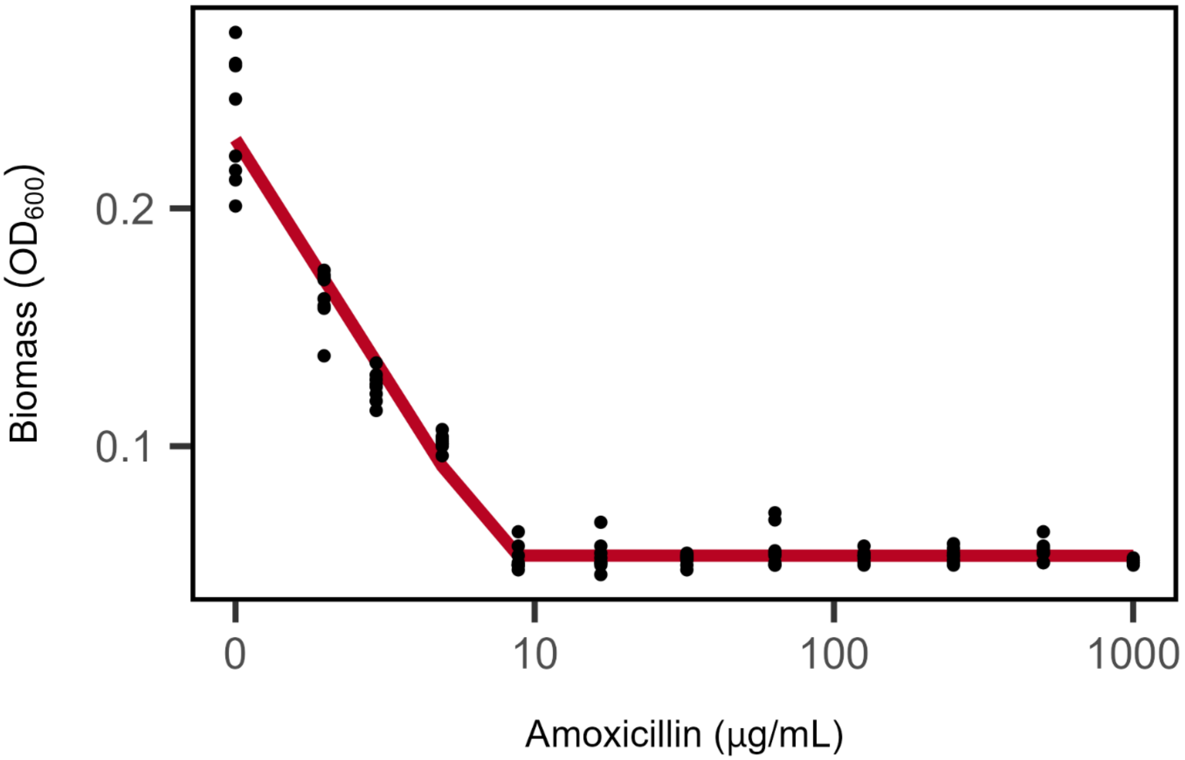

**Fig. S2.**

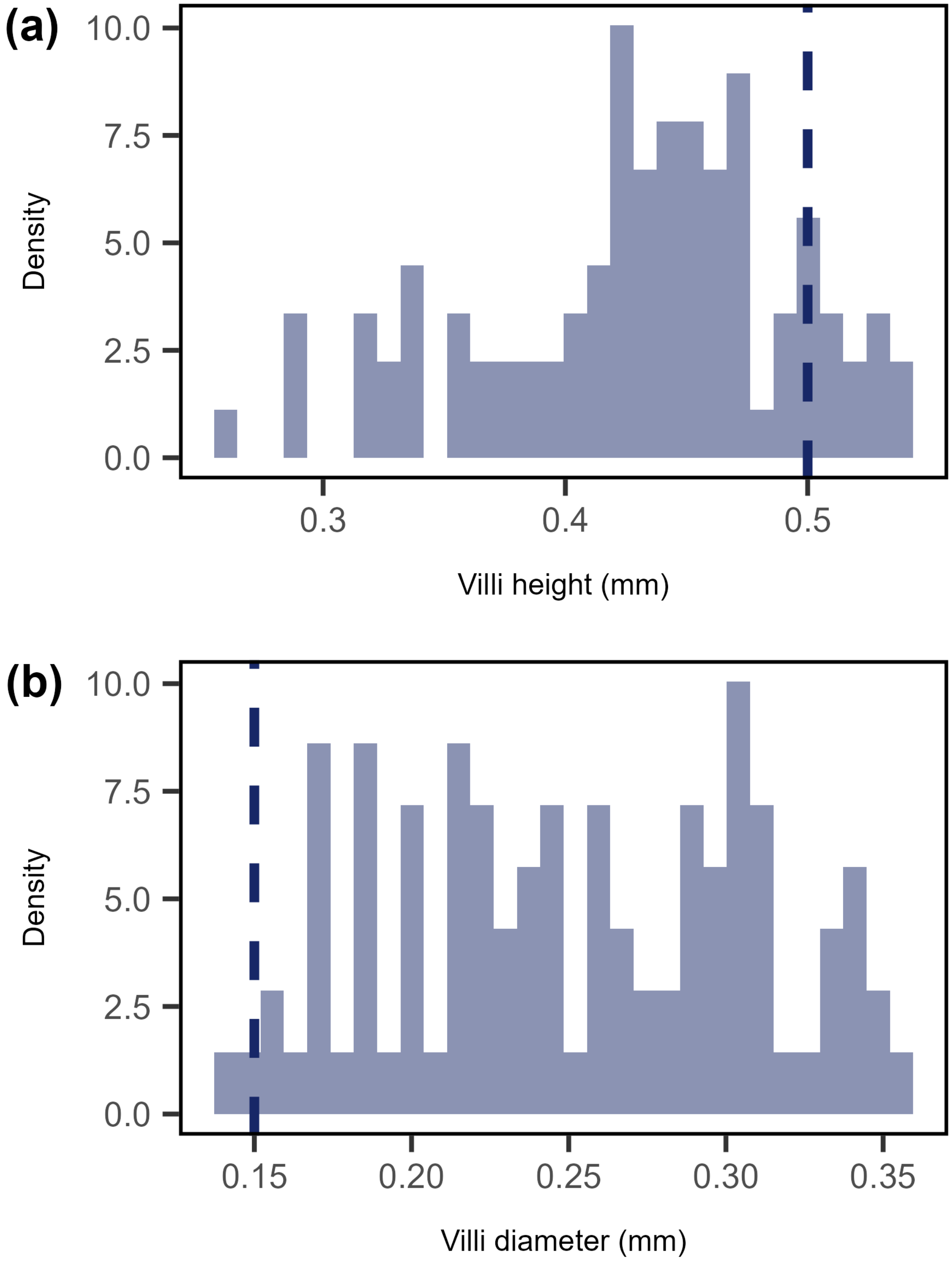
(a)
10.0
7.5
5.0
2.5
0.0
Density
0.3
0.4
0.5
Villi height (mm)
(b)
10.0
7.5
5.0
2.5
0.0
Density
0.15
0.20
0.25
0.30
0.35
Villi diameter (mm)

**Fig. S3.**

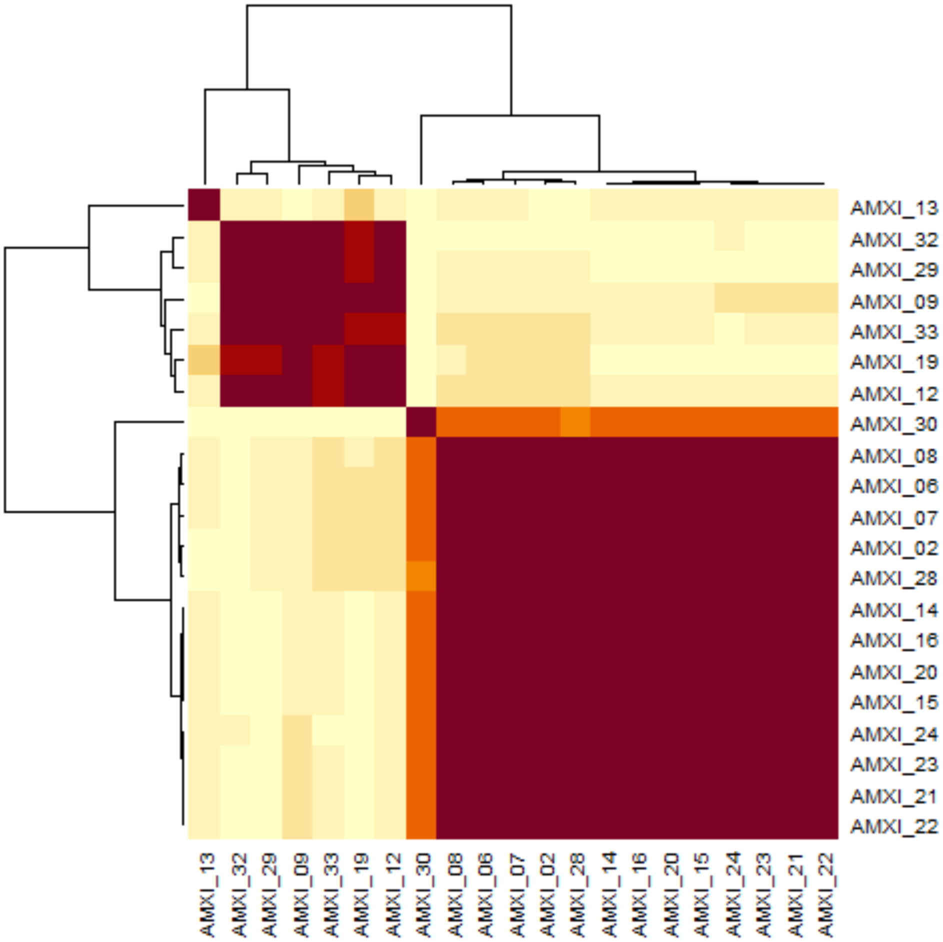
AMXI_13
AMXI_32
AMXI_29
AMXI_09
AMXI_33
AMXI_19
AMXI_12
AMXI_30
AMXI_08
AMXI_06
AMXI_07
AMXI_02
AMXI_28
AMXI_14
AMXI_16
AMXI_20
AMXI_15
AMXI_24
AMXI_23
AMXI_21
AMXI_22
AMXI_13
AMXI_32
AMXI_29
AMXI_09
AMXI_33
AMXI_19
AMXI_12
AMXI_30
AMXI_08
AMXI_06
AMXI_07
AMXI_02
AMXI_28
AMXI_14
AMXI_16
AMXI_20
AMXI_15
AMXI_24
AMXI_23
AMXI_21
AMXI_22
